# Deciphering defective subventricular adult neurogenesis in cyclin D2-deficient mice

**DOI:** 10.1101/419382

**Authors:** Rafał Płatek, Leszek Kaczmarek, Artur Czupryn

## Abstract

Adult neurogenesis occurring in the brain of adult mammals is considered to have potential therapeutical applications. New neurons are produced constitutively from postnatal neural stem cells/precursors residing in two neurogenic regions: the subventricular zone (SVZ) of the lateral ventricles and the subgranular layer in the dentate gyrus of the hippocampus. Newly-generated neuroblasts from the SVZ migrate long distance towards the olfactory bulb and repopulate different subtypes of inhibitory interneurons modulating the olfactory processing. It was reported that cyclin D2 knockout mice (cD2-KO) present reduced generation of new hippocampal neurons, however proliferation deficiency and mechanisms responsible for dysregulation of subventricular precursors, derived progenitors, and olfactory interneurons need to be detaily investigated. In this report, proliferative activity of different subpopulations of SVZ neural precursors, cell migration, and differentiation in cD2-KO mice was characterized. For that goal, EdU, a thymidine analogue, proliferation mapping combined with multi-epitope immunohistochemical detection of endogenous stage-specific cell markers was carried out. Severely reduced number of newly-generated cells in the subventricular niche was demonstrated that was not accompanied by increased level of apoptotic death. Surprisingly, the number of B1 quiescent precursor subpopulation was not affected, whereas the number of B1 type active primary precursors, intermediate/transiently-amplifying progenitors (C type cells), and neuroblasts (A type cells) were reduced. The analyses suggest that cycline D2 might be critical for transition of B1 precursor quiescent cells into B1 active cells. We also demonstrate that the subpopulation of calbindin interneurons is reduced in the olfactory bulb. Deciphering processes underlying a potential modulation of intensity of adult neurogenesis at the cellular levels could lead to replacement therapies after injury, stroke, or neurodegenerative disease in the central nervous system.

## INTRODUCTION

Most adult mammals are capable of producing new neurons (Altman, 1963; Eriksson et al., 1998). Adult neurogenesis, proliferation and differentiation of neural progenitors, takes place in specific neurogenic brain regions, which are the subgranular zone (SGZ) of the hippocampal dentate gyrus (DG) and the subventricular zone (SVZ) also referred to as the subependymal zone (SEZ) along the lateral ventricles (LVs) (Altman and Das, 1965; Luskin 1993; Morshead et al., 1994; Eriksson et al., 1998). SGZ and SVZ neurogenic niches produce DG neurons and OB interneurons, respectively throughout the lifespan (Luskin 1993; Cameron et al., 1993; Gage, 2000; Merkle et al., 2007). Adult neurogenesis is of great attention due to proposed contribution to learning and memory phenomena and potential role in repopulating injured CNS regions after neurotrauma events (Arruda-Carvalho et al, 2011, Arlotta et al., 2003, Arvidsson et al, 2002, Toad and Gage, 2018). However, adult neurogenesis functional significance for the cognitive processes and knowledge of its regulatory mechanisms for restorative purposes is still insufficient.

Adult neurogenesis requires instant proliferation of neural precursors. Among molecules controlling cell division are cyclins D - cell cycle regulatory proteins, that act as regulatory subunits in complexes with cyclin-dependent kinases (CDKs) (Sherr and Roberts, 1999). There are three D type cyclins: D1, D2 and D3, with partially overlapping tissue expression pattern.

Neural stem cells from brain neurogenic niches also depend on cyclin D in requirements for proliferation (Ciemerych et al., 2002; Kowalczyk et al., 2004; Jaholkowski et al., 2009). Kowalczyk et al. (2004) reported significance of cyclin D2 in particular, showing that in an adult wild type (WT) mice it is the only form expressed out of D1, D2 and D3 and selective deficiency of cyclin D2, but not D1, resulted in impaired neurogenesis. Cyclin D2-KO (cD2-KO) mice were almost completely deprived of BrdU(+) cells both in hippocampal DG and in the OB. None of the single BrdU(+) cells in hippocampi of D2-KO mice co-localized with TuJ1 or NeuN neuronal markers, in contrast to WT littermates. In line are the results that cyclin D2 deficient mice were unable to upregulate hippocampal neurogenesis after induction by exposure to enriched environment although retained ability to upregulate astrocyte proliferation after cortex lesioning (Kowalczyk et al., 2004).

Several studies addressed the role of adult neurogenesis in learning and memory processes in rodents. Cyclin D2 knockout mice used as a model for virtually missing adult brain neurogenesis showed that almost complete lack of progenitor proliferation in SGZ, does not influence fear and spatial learning and memory (Jaholkowski et al., 2009; Urbach et al., 2012). cD2-KO mice although presenting 80-90% reduction in number of BrdU(+) cells in granular layer (GL) of DG, showed normal performance in sensorimotor tasks and in procedural learning. No deficits in context, cue and context-remote fear conditioning were reported and D2 deficient mice performed significantly better in trace-cue fear conditioning compared to WT littermates (Jaholkowski et al., 2009). Morris water maze test, IntelliCage system (Jaholkowski et al., 2009) and Barnes maze (Urbach et al., 2012) proved additionally that cD2-KO mice also had no marked deficits in spatial learning and memory tests.

Little is known about influence of cyclin D2-defficiency on neurogenesis in the subventricular zone. Kowalczyk et al. (2004) reported changed proliferative general properties of the SVZ neurogenic niche, however no spatial nor populational analyses were performed. What is important, cD2-KO mice showed quite severe olfactory dysfunctions, as they were unable to find hidden food under the cage bedding (Jaholkowski et al., 2009). This suggests an important role of constant addition of SVZ-derived newly-generated neurons into the olfactory bulb network for proper function and smell discrimination.

The goal of our work was to investigate in details proliferative activity of different subpopulations of SVZ neural precursors, cell migration, and differentiation of target interneurons of the olfactory bulb in cD2-KO mice. To fulfill these aims, we have decided to use a new generation thymidine analogue—EdU, for proliferation mapping which we combined with multi-epitope immunohistochemical detection of endogenous stage-specific cell markers of different subpopulations of precusors/progenitors, neuroblasts, and interneurons in the SVZ and the olfactory bulb in cD2-KO mice.

## MATERIALS AND METHODS

### Animals

Mouse line with inactive gene of Ccnd2 responsible for expression of cyclin D2 was generated by group of Dr. P. Sicinski (Sicinski et al, 1996), and was backcrossed into C57BL/6 strain background over 20 times in our group. Cyclin D2 heterozygotes were used as breeding pairs at the animal facility of the Nencki Institute in Warsaw and their cyclin D2 -/- (Cyclin D2 knockout, cD2-KO) and +/+ (wildtype, WT) mice, 3-6-month-old, males and females were used in all the experiments. Animals were given free access to water and pellet food and were housed under standard humidity and temperature conditions on a 12 h light/dark cycle. Experimental protocols involving animals and care were approved by the First Local Ethics Committee in Warsaw (permit No. 343/2012) and strictly followed the rules of the Polish Animal Protection Act. Animal experimentation was carried out in accordance with EU directive 2010/63/EU.

### EdU injections

Animals were injected with EdU-a thymidine analogue (Thermo Fisher Scientific), 50mg/kg in a single dose i.p., twice a day (8-10h apart) for 5 days (10 times in total) and subsequently sacrified with fixative perfusion 48-60h after the last EdU injection.

### Tissue preparation

Two days after the last EdU injection mice were euthanized with a lethal dose of Morbital (80 mg/kg) or with the mixture of Medetomidine (1 mg/kg) and Ketamine (75 mg/kg) i.p. For fluorescence analysis, mice were perfused transcardially with 0.1 M perfusion buffer (PBS), pH 7.4 (100 mM phosphate buffer; 140 mM NaCl; 3.4 mM KCl; 6 mM NaHCO3) followed by 4% paraformaldehyde in perfusion buffer. Next, the brains were removed from the skulls and postfixed in the same fixative overnight at 4°C. Tissue was then cryoprotected stepwise in 10%, 20% and 30% sucrose in 0.01 M PB at 4<u>°</u>C, then cut on 20-30μm thick free-floating or glass mounted sections using Leica cryostat.

### Immunofluorescence

Prior to immunolabeling sections were washed 3 times 5 min each with 0.01 M PBS pH 7.4 or 20mM Tris-buffered saline buffer (TBS) pH 7.4 and blocked and permeabilized for 30 min with 5% normal donkey serum (NDS) in PBST or TBST respectively (PBS or TBS + 0.2% Triton X-100). In case of caspase 3 protocol, before the blocking step, sections were first immersed in 1% SDS for 5 min at room temperature, then washed 3 times for 5 min and next blocked in NDS. Next, for protein antigen detection sections were incubated overnight at 4°C with the following primary antibodies directed specifically against the following epitopes: Ki67 (1:1500; Abcam ab16667), cleaved caspase 3 (1:150; R&D AF835), doublecortin (1:500; Abcam ab18723), GFAP (1:200; Sigma), EGFR (1:300; Santa Cruz Biotechnology sc-03), Calbindin D-28k (1:1500; Swant CB300), Calretinin (1:1500; Swant 6B33), tyrosine hydroxylase (1:300; Abcam ab137869). Sections were on the next day washed 3 times 5 min each with 0.01 M PBS pH 7.4 or 20 mM TBS pH 7.4, then incubated with F(ab)2 fragments of secondary IgG antibodies, all raised in donkey against mouse, rabbit, or goat and conjugated with Alexa Fluor-405, −488, −594 or −647 (Jackson Immunoreseach). As a positive control for detection of apoptotic cells during immunohistochemical visualization of caspase 3 (validation of applied antibody and experimental approach), we used similar brain sections obtained from WT animals after middle cerebral artery occlusion (MCAO) (kind gift of representative sections from W. Karunakaran from our Institute, for procedure see reference Zawadzka et al. 2012),

### EdU detection

EdU was visualised using Click-iT chemistry with Click-iT Cell Reaction Buffer Kit (Thermo Fisher Scientific #C10269). If EdU labeling was combined with immunostaining, Edu detection was performed first. Briefly, free-floating sections were washed with 20 mM TBS pH 7.4 three times for five min, followed by incubation in TBST for 30 min for permabilization and washed once afterwards in TBS. Next sections were incubated in an EdU detecting mixture (500μl/well) containing: 1) Reaction buffer, 2) CuSO4 (1:50), 3) Buffer additive (1:10) and 4) 3 μM Alexa Fluor 594 azide for 30 min room temperature (mixture prepared within 15 min before usage) according to the manufacturer’s protocol. For negative control of labeling specificity, CuSO4 was omitted, were copper catalyzes triazole formation from an alkyne (EdU-alkyne) and an azide (Alexa Fluor 594-zide). Next sections were washed in TBS (3×5min), stained with Hoechst 33342, washed in TBS and coverslipped for analysis. If Edu detection was combined with immunostaining, after labeling with EdU reaction cocktail, sections were washed in TBS and followed with blocking in serum and incubation with primary antibodies and processed as described above.

### TUNEL staining

For analysis of apoptosis level in the SVZ, TUNEL method was applied (In Situ Cell Death Detection Kit, Fluorescein; Roche #11684795910), which enables labeling of apoptotic cells upon detection of DNA strand brakes and labeling the free 3’-OH groups with fluorescein modified nucleotides. The 18-20 μm thick coronal sections mounted on the slides were used. Briefly, sections were washed in 0.1M PBS pH 7.4 three times for 5 min and incubated in PBS + 0.5% Triton X-100 in 85°C for 20min, then cooled and washed in PBS (3×5min). Next positive control sections were incubated in DNase I (300 U/ml) in PBS for 10min while experimental sections in PBS. Then all sections were incubated in TUNEL reaction mixture (50 μl/slide) containing: 1) label solution and 2) enzyme solution (1:10) for 1h at 37°C. For negative control of labeling specificity enzyme solution containing terminal deoxynucleotidyl transferase (TdT) was omitted which catalyzes polymerization of deoxyribonucleotides to the 3’-end of single- and double-stranded DNA. Next sections were washed in PBS + 0.5% Triton X-100 (3×5min), and coverslipped with Fluoromount G with DAPI for analysis.

### Analysis of proliferation activity in SVZ

For analysis of proliferation activity in the SVZ, 30 μm thick coronal sections of the brains from cyclin D2 WT (n=3) and KO (n=3) mice were used, starting with the SVZ sections containing clearly distinguishable three distinct walls - lateral, dorsal and medial and ending before the appearance of the dorsal part of the hippocampus. Seven to eight sections along anterior-posterior SVZ axis were divided into the rostral (five sections) and caudal (two-three sections) SVZ. After detection of EdU, sections were scanned using confocal microscope Leica SP5 and pictures of SVZs were taken using 10x objective from 19-20 μm depth. In sections from the rostral SVZ five areas were distinguished: 1) lateral wall (L), dorsal wall (D), medial wall (M), top cone (T), and bottom cone (B) (see also the Results section). In sections from the caudal SVZ three areas were distinguished: 1) L, 2) D, and 3) T. For the analysis z-stacks were flattened to one plane to count all EdU positive profiles in the 50 μm wide area from the ventricle wall in case of L, D, and M, and within the distance of 100 μm from the ventricle wall in the T and B regions.

### Analysis of phenotypes of progenitors in the SVZ

For phenotypic analysis of cell classes in the SVZ, 30 μm-thick coronal sections of the brains from cyclin D2 WT (n=4) and KO (n=3) mice were used. First five-six sections from rostral SVZ (from analysis of proliferation activity) were chosen. Sections were subjected to EdU detection with Alexa Fluor 594-azide, and GFAP and EGFR immunodetection with donkey anti-mouse Alexa Fluor 488 and donkey anti-rabbit Alexa Fluor 647, respectively, and Hoechst 33342 labeling according to the protocols above. Next sections were scanned using Spinning Disc confocal microscope (Zeiss Inverted Axio Observer Z.1) and photographs of lateral walls from the SVZs were taken under 40x oil objective from 30 μm depth with 0.3μm z-step. We distinguished 3 classes of cells in the SVZ: 1) quiescent B1 cell type [NSCs, B1q]: GFAP(+), EGFR(-), EdU(-), Hoechst(+); 2) active B1 cell type [B1a]: GFAP(+), EGFR(+), EdU(+) or EdU (-), Hoechst(+); and 3) C cell type [intermediate progenitor cells, IPCs; transit amplifying precursors TaPs] together with A cell type [neuroblasts]: GFAP(-), EGFR(+), EdU(+) or EdU(-), Hoechst(+). Final quantifications were performed on the middle z-stacks from 20 μm in-depth and specific classes of cells were recognized in the 50 μm wide area from the ventricle wall.

### Statistical analysis

For statistical comparison of differences between independent samples, Student’s t-test was used.

## RESULTS AND DISCUSSION

First, we aimed to investigate in details whether cell divisions were affected in all or only selected walls (anatomical and functional sub-domains) of the subventricular zone around the lateral ventricles in adult mice with defective adult neurogenesis due to deactivation of the cyclin D2 expression.

In order to investigate cell proliferation activity, we used a method of EdU incorporation, a modified method of bromodeoxyuridine (BrdU) cell mapping. EdU as well as BrdU, both thymidine analogs, after systemic administration to experimental animals, are incorporated to DNA during its replication in the S phase of the cell division cycle. The proliferative activity was studied in the anterio-posterior axis as well as in the dorso-ventral axis of the subventricular zone in a series of 30-µm-thick coronal sections of brains from cD2-KO and WT mice.

Our preliminary analyses of EdU detection in brain coronal sections of adult cD2-KO and WT mice demonstrated that proliferation intensity differs in the rostral and caudal compartments of the SVZ. For that reason, we have carried out the analyses of EdU-positive cells separately in these two general compartments. The border between rostral and caudal parts of the SVZ was arbitrary considered at a coronal section in which the most rostral hippocampal structures become distinguished. Representative examples of the rostral and caudal transections of the SVZ are presented on Fig. 1A and Fig 1B. Moreover, we have adjusted our experimental approach of cell counting in regards to different tendencies of localization of EdU-positive cells in these two SVZ areas. In the rostral compartment, the number of dividing cells was calculated in all three walls of the SVZ: dorsal, medial, and lateral, while in the caudal compartment within only two walls: lateral and dorsal, and the neurogenic niche in all cases was defined strictly within 50-µm width along the SVZ. Additionally, in both compartments, we clearly distinguished characteristic cohort of proliferating cells at the crossing of the dorso-lateral walls, lineally-organized (and semi-laterally directed) group of EdU(+) cells within the arbitrary counting distance of 100 µm from the SVZ, called by us the top cone (T) on the Fig. 1A. Only in the rostral SVZ compartment, a characteristic cohort of EdU(+) cells was also visible at the junction of medio-lateral walls, an accumulative area of EdU(+) cells within the arbitrary counting distance of 100 µm, called by us the bottom cone (B) on the Fig. 1A.

**Fig. 1.**
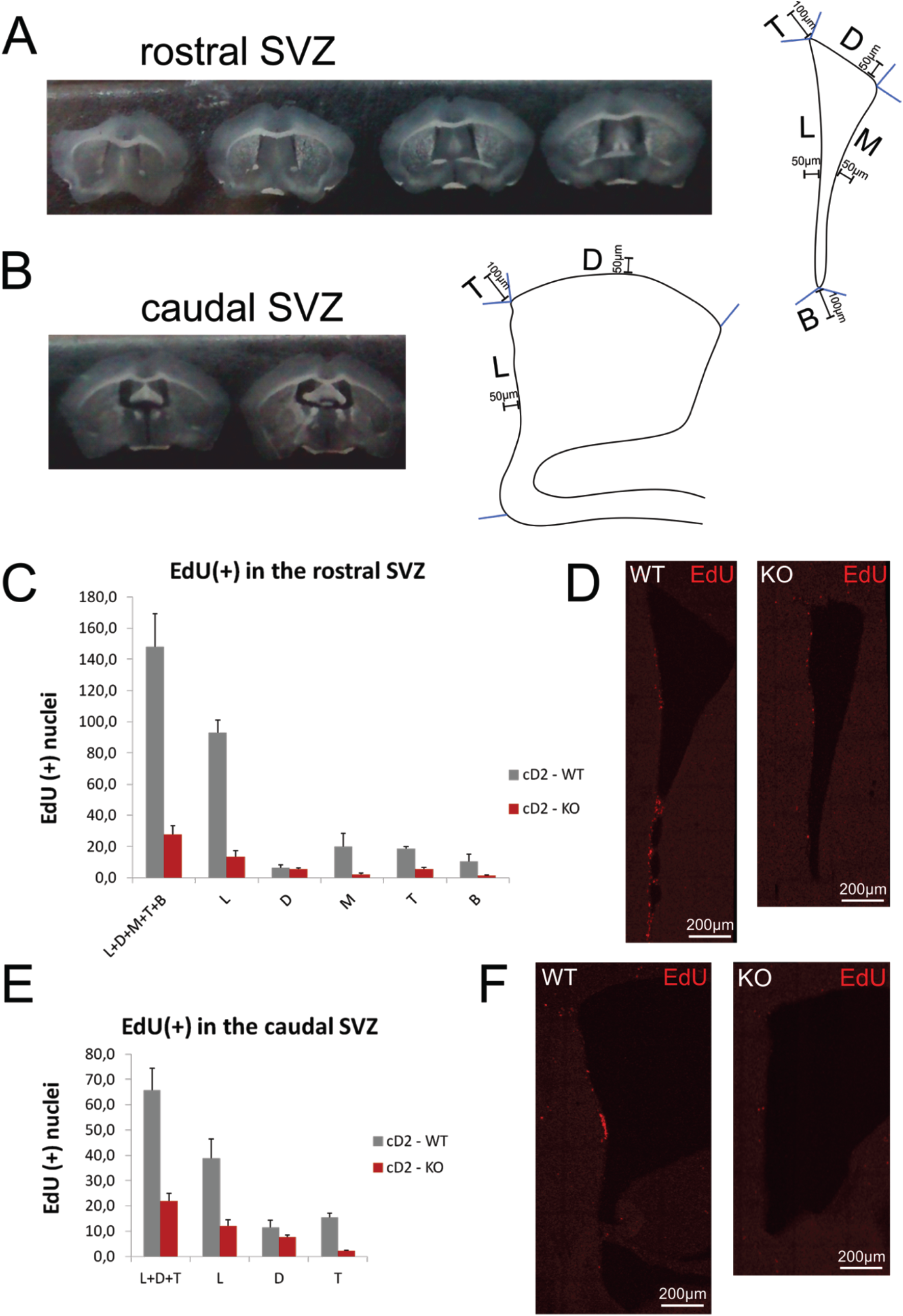
Cyclin D2-KO mice show significantly reduced proliferation activity in the SVZ. (A-B) Photographs of representative coronal brain sections containing the rostral (A) and caudal (B) SVZ along anterio-posterior brain axis chosen for analysis. Schematic drawings of the compartments analyzed in the rostral (A) and caudal (B) SVZ. (C-D) A graph and microphotographs show reduced number of proliferation marker positive(EdU+) profiles in majority of the rostral SVZ compartments in cD2-KO mice compared to WT animals. (E-F) Graph and microphotographs show reduced number of EdU(+) profiles in the caudal SVZ of cD2-KO mice.

A quantitative analysis of the distribution of dividing cells revealed significant global decrease in cell proliferation in the SVZ neurogenic niche of the cyclin D2-defficient mice (Fig. 1C-D). Detailed analysis demonstrated that cell proliferation in cD2-KO mice was significantly affected in all anatomical sub-regions (walls and cones) distinguished as sub-areas for the counting, only the dorsal wall of the SVZ seemed to be less affected (both in the rostral and caudal SVZ). The most prominent decreases of irrelative values in cell proliferation was observed in the lateral wall, the medial wall, and in the laterally-oriented top cone (values on Fig 1C, 1D). These observations support a notion that deactivation of cyclin D2 produce decrease in cell proliferation along the dorso-ventral axis (cD2-KO vs. WT differences in walls and cones within a coronal transection of the SVZ) and also along the anterio-posterior axis (relative differences between rostral and caudal SVZ compartments). Taking together, we confirm general substantial changes in proliferation and provide detailed subregional measurements in the neurogenic niche of the SVZ of adult mice with cyclin D2 deficiency.

We have addressed one of potential hypothesis that the decrease in the number of newly-generated cells might be caused by an elevated cell death occurring in the subventricular neurogenic niche of cD2-KO mice. In order to quantify the progress of programmed cell death, we performed analysis of apoptosis level measured by the presence of activated (cleaved) caspase 3, a classical hallmark of apoptotic death, which is present in the medial and final stage of dying cells. Using immunohistochemical detection of activated caspase 3, we observed no cells either in cD2-KO or WT mice. In order to validate that applied antibody and experimental approach specifically and effectively detects apoptotic cells, we have used a positive control tissue. For that purpose, we used similar brain sections obtained from animals after middle cerebral artery occlusion (MCAO) (for reference see the section *Material and Methods*, and Zawadzka et al. 2012), a classic model of brain ischemia, which is widely reported in the literature as being followed with robust apoptotic cell death in the infarct area. We processed the MCAO brain sections simultaneously during immunohistochemical cleaved caspase 3 detection together with a series of sections from cD2-KO and WT mice. Epifluorescence microscopic analysis revealed immunopositive cells in the tissue after MCAO but no labelled cells in the SVZ in the brains of cD2-KO and WT mice (Fig. 2A). This can be concluded that SVZ and the neighboring areas in the cD2-KO mice do not contain increased level of apoptosis in comparison to untreated WT mice.

**Fig. 2.**
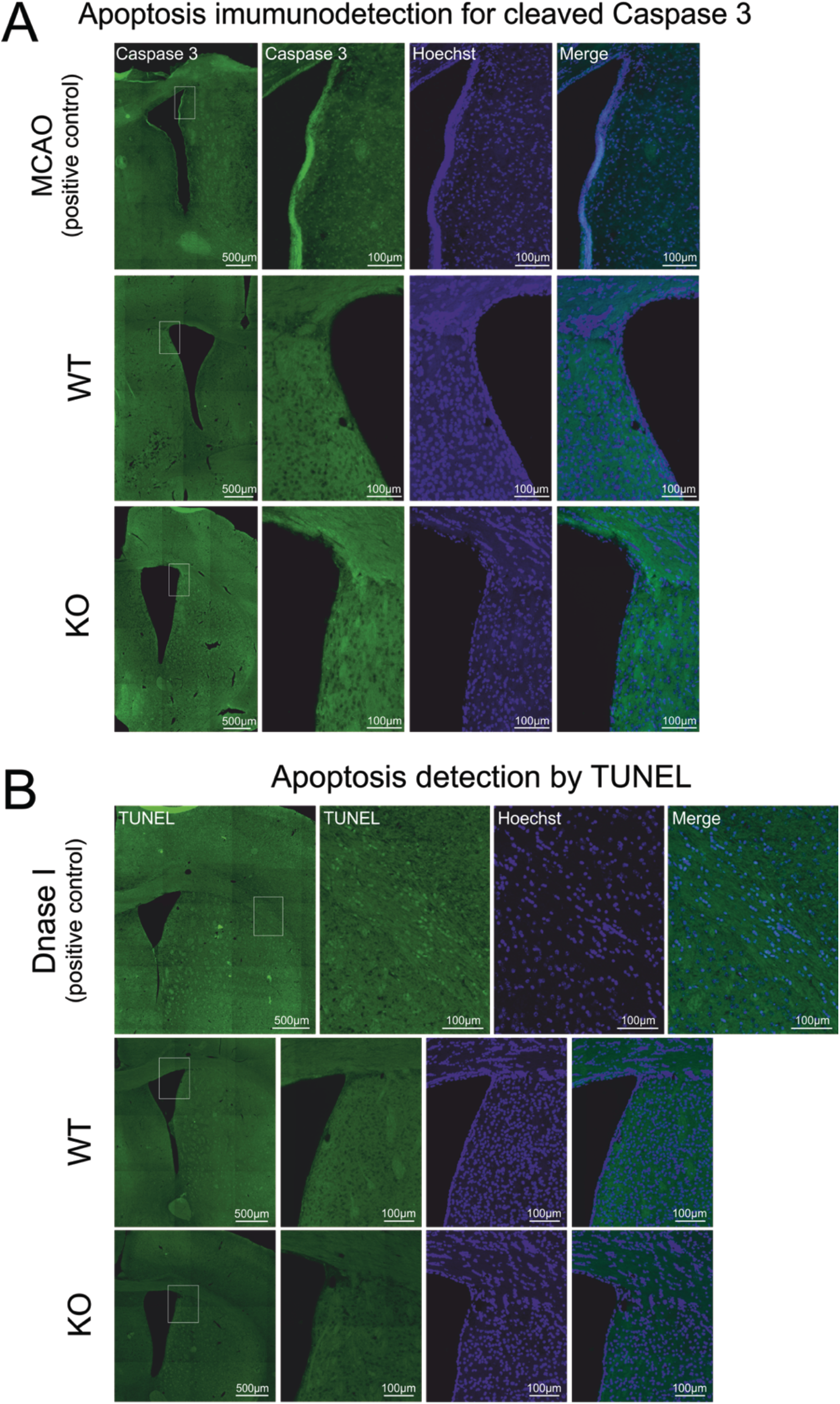
Upregulation of apoptosis does not contribute to reduced number of actively dividing cells in the SVZ of cD2-KO mice. (A) Immunostaining for cleaved caspase 3. There is no positive signal of active caspase 3 in the SVZ of cD2-WT and cD2-KO mice under physiological conditions. (B) Detection of apoptosis using TUNEL. There is no positive signal for DNA strand brakes in the SVZ of cD2-WT and cD2-KO mice under physiological conditions.

In order to confirm these observations with reliable alternative approach of apoptosis detection, we applied TUNEL staining. This method detects chromosomal DNA fragmentation in the double-strand DNA breaks as characteristic structural changes during the programmed cell death. The TUNEL reaction in brain sections from cD2-KO and WT mice revealed no or very sporadic apoptotic cells within the SVZ, in all three walls of the lateral ventricles, not only in the width of the neurogenic zone around the lateral ventricles (50µm-wide area along the SVZ walls or 100µm-wide area in the top and bottom cones) but also in potentially defectively migrating cells in the distance from the SVZ (Fig. 2B). Positive controls— sections treated with DNase I—presented successful confirmation of the applied method. Taking together, the results obtained from the two approaches for apoptosis detection have rule out a hypothesis that the decrease in number of newly generated cells in the subventricular neurogenic niche in the cD2-KO mice is caused by death of post-mitotic cells. Summing up all results presented up to this point, we can conclude that cyclin D2-deficient mice show lower level of cell proliferation in the SVZ, and this phenomenon occurs without an influence of cell apoptosis.

In order to investigate what type or types of cells are directly affected by lowered proliferation level, we have characterized cell populations of the subventricular neurogenic niche in the cD2-KO mice and compared to the composition of cells in WT mice. At first, we observed the lowered number of neural stem cells (precursors) and intensively proliferating secondary progenitors, both immunoreactive for the epidermal growth factor receptor (EGFR, Fig. 3A). This protein marker is primarily expressed on a B1 cells (primary postnatal precursors residing in the SVZ, including both B1 quiescent and B1 slowly dividing precursors) and C cells (intermediate progenitor cells, secondarily derived from B1 cells). Also, we detected significant defects in the number of migrating neuroblasts immunopositive for doublecortin (Dcx) (Fig. 3B).

**Fig.3.**
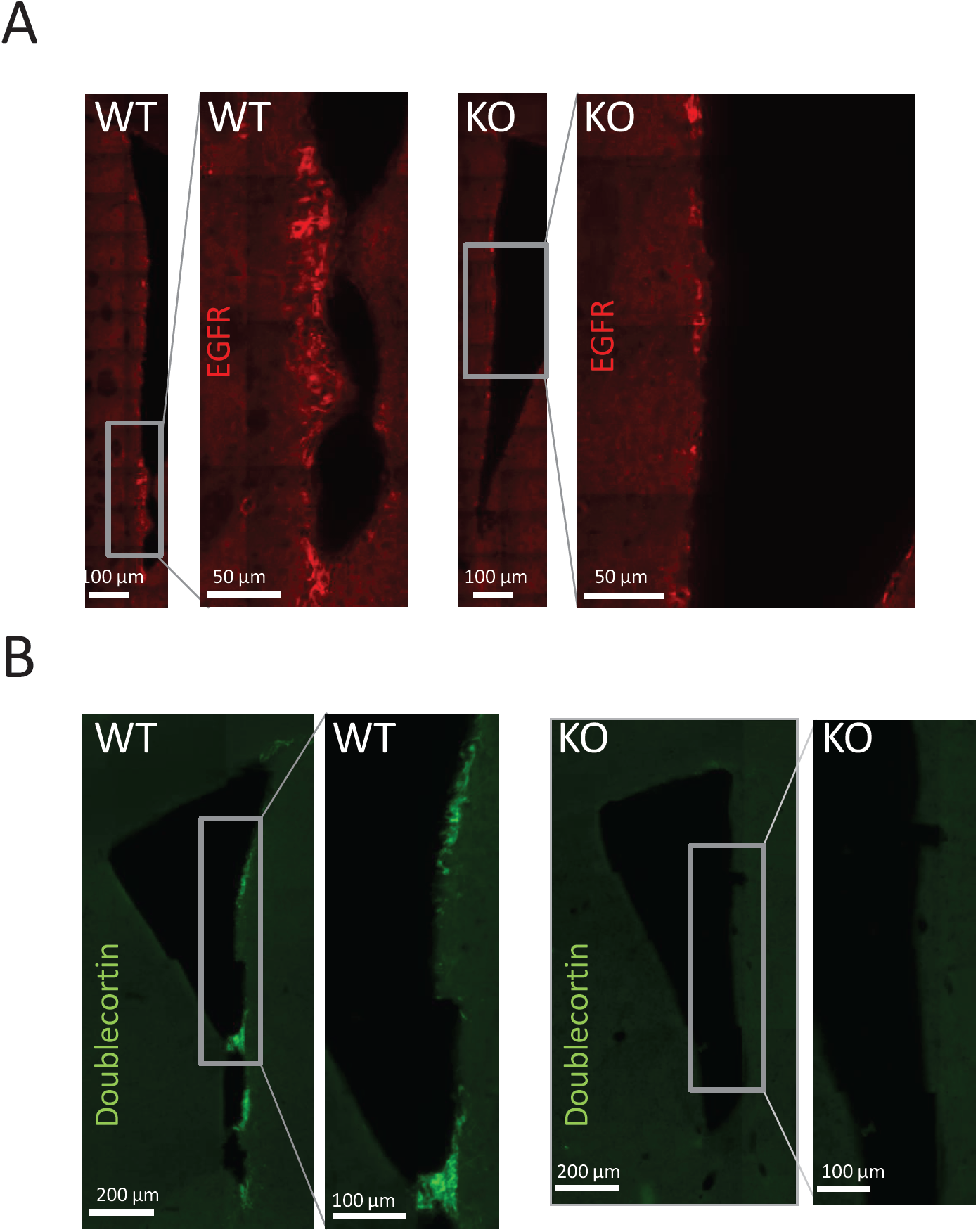
Number of progenitors and neurblasts is downregulated in the SVZ of cD2-KO. (A) Immunostaining for epidermal growth factor receptor (EGFR). Microphotographs show fewer EGFR(+) B1 active and C type cells (TaPs) in the lateral wall of cD2-KO vs cD2-WT mice. (B) Immunostaining for doublecortin (Dcx). Microphotographs show that the lateral wall of the SVZ from cD2-KO mice is largely depleted of neuroblasts compared to WT animals.

These observations provoked us to perform more comprehensive quantitative analyses of cell populations. Because there are no single cell markers specific for each subpopulation of cell subclasses in the subventricular niche, they were identified in our analyses by simultaneous detection of multiple epitopes. Additionally, we combined this multi-epitope-immunoreaction with Click-iT-chemistry reaction for EdU visualization for dividing/proliferating cells. A combination of identifying cell markers was as described below. Neural precursors (B1 quiescent cell type) quiescent with cell proliferation properties were identified as: GFAP(+), EGFR(-), EdU(-). Activated neural precursors (B1 activated cell type), with their unique long cell cycle have divided or not during a limited period of experimental EdU delivery to mice, are: GFAP(+), EGFR(+), both EdU(+) and EdU(-). Originating from them intensively dividing cells known as progenitors (C cell type, known also as intermediate progenitor cell, IPS, or transiently-amplifying progenitor, TaP) have an immunohistochemical phenotype of: GFAP(-), EGFR(+), mostly EdU(+) but also less frequently EdU(-), and the same phenotype was presented by neuroblasts (A cell type), mostly EdU(-). In order to unambiguously identify separate cells and provide reliable cell counting, cell nuclei were in all cases counterstained with DNA-specific dye Hoechst 33342. Representative photomicrographs of phenotype subclasses are shown on Fig. 4A. As reported by us above, the highest number of EdU(+) proliferating cells both in WT and cD2-KO mice was in the neurogenic zone of the lateral wall (i.e. this wall in the major site for the subventricular adult neurogenesis). For that reason, we performed multiple epitope cell counting and complex discrimination to the listed subpopulation only in the lateral wall of the subventricular zone (strictly within 50-µm wide distance from the ventricles, compare Fig. 1).

**Fig.4.**
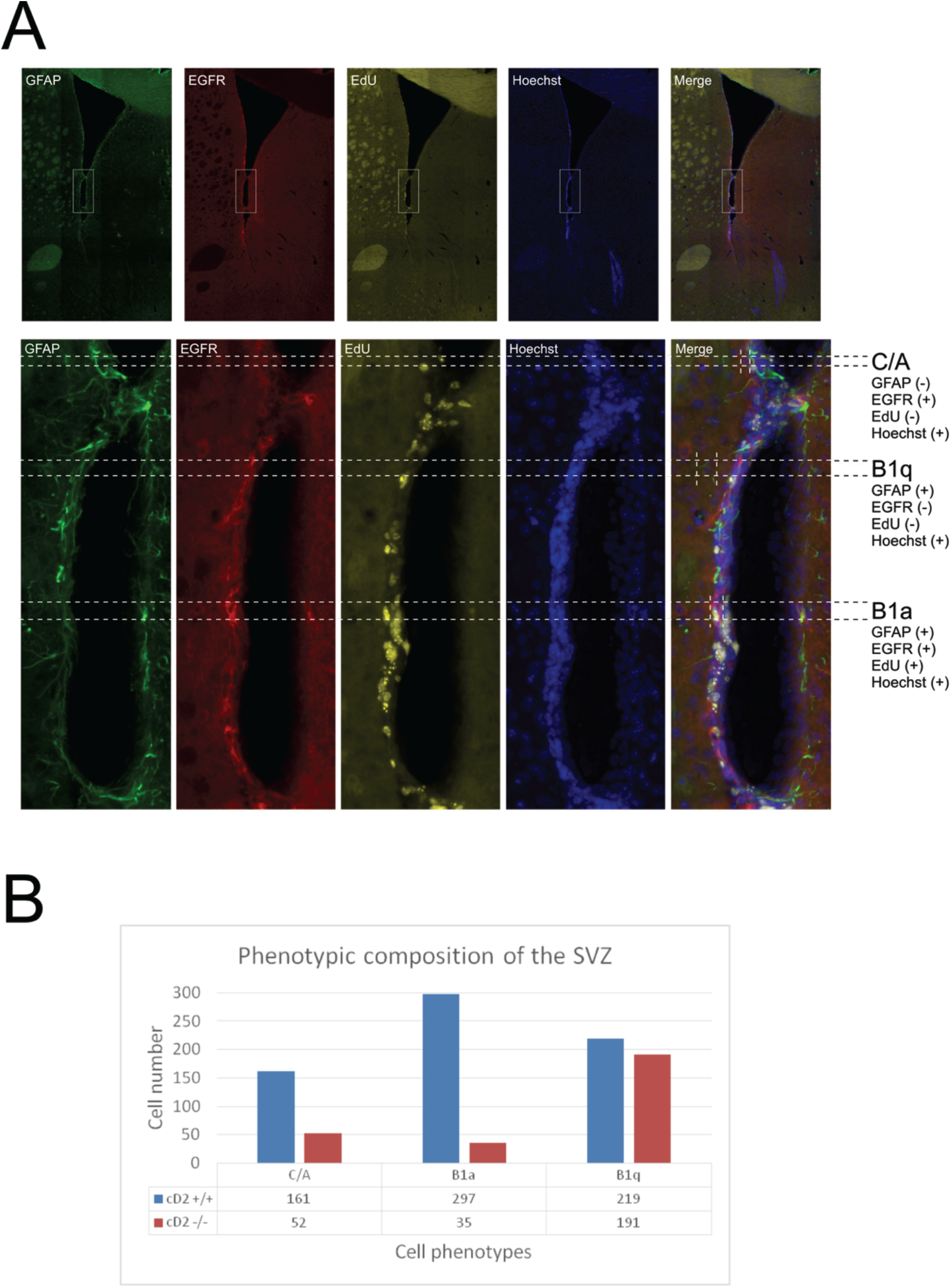
Cyclin D2 deficiency affects transition from B1 quiescent to B1 active progenitors. (A) Microphotographs of representative SVZ from cD2-WTmice showing immunostaining for GFAP (green), EGFR (red), EdU (yellow) and Hoechst (blue) for identification of four classes of cells in the SVZ neurogenic niche: 1) B1 quiescent precursors (B1q): GFAP(+), EGFR(-), EdU(-), Hoechst(+); 2) B1 active precursors (B1a): GFAP(+), EGFR(+), EdU(+) or EdU(-), Hoechst(+) and 3) intermediateprogenitor cells/transit amplifying progenitors (IPCs/TaPs) and neuroblasts: GFAP(-), EGFR(+), EdU(+) or EdU(-), Hoechst(+). (B) Analysis of cellular composition in the SVZ lateral wall indicates that lack of cyclin D2 influences transition from B1q cells to B1a progenitors.

After careful manual discrimination of cells in representative sections into described above pools of cells, we calculated that number of B1 quiescent cells were, surprisingly, unchanged in cD2-KO *vs*. control neurogenic niche in WT mice (Fig. 4B).

On the other hand, the number of B1 active cells and the collective number of intermediate/transiently-amplifying progenitors and neuroblasts were significantly decreased in cyclin D2-deficient mice (Fig. 4B). Thus, these analyses can be concluded that cycline D2 might be critical for transition of B1 precursor quiescent cells into active B1 rarely dividing cells.

Finally, it was substantial to investigate whether changed numbers of proliferating cells in the subventricular neurogenic niches of adult cD2-KO mice might reflect differences in the composition or spatial distribution of subpopulations of inhibitory interneurons in the olfactory bulb originating in a substantial pool from the postnatal SVZ precursors.

We have noticed, that the brains of cD2-KO mice were smaller than in WT mice, especially the olfactory bulbs, as reported by Kowalczyk et al., (2004). Moreover, applying histochemical Nissl staining, we demonstrated that gross morphology of cell layers (thickness of layers and cell density) within the olfactory bulbs was significantly changed (Fig. 5).

**Fig.5.**
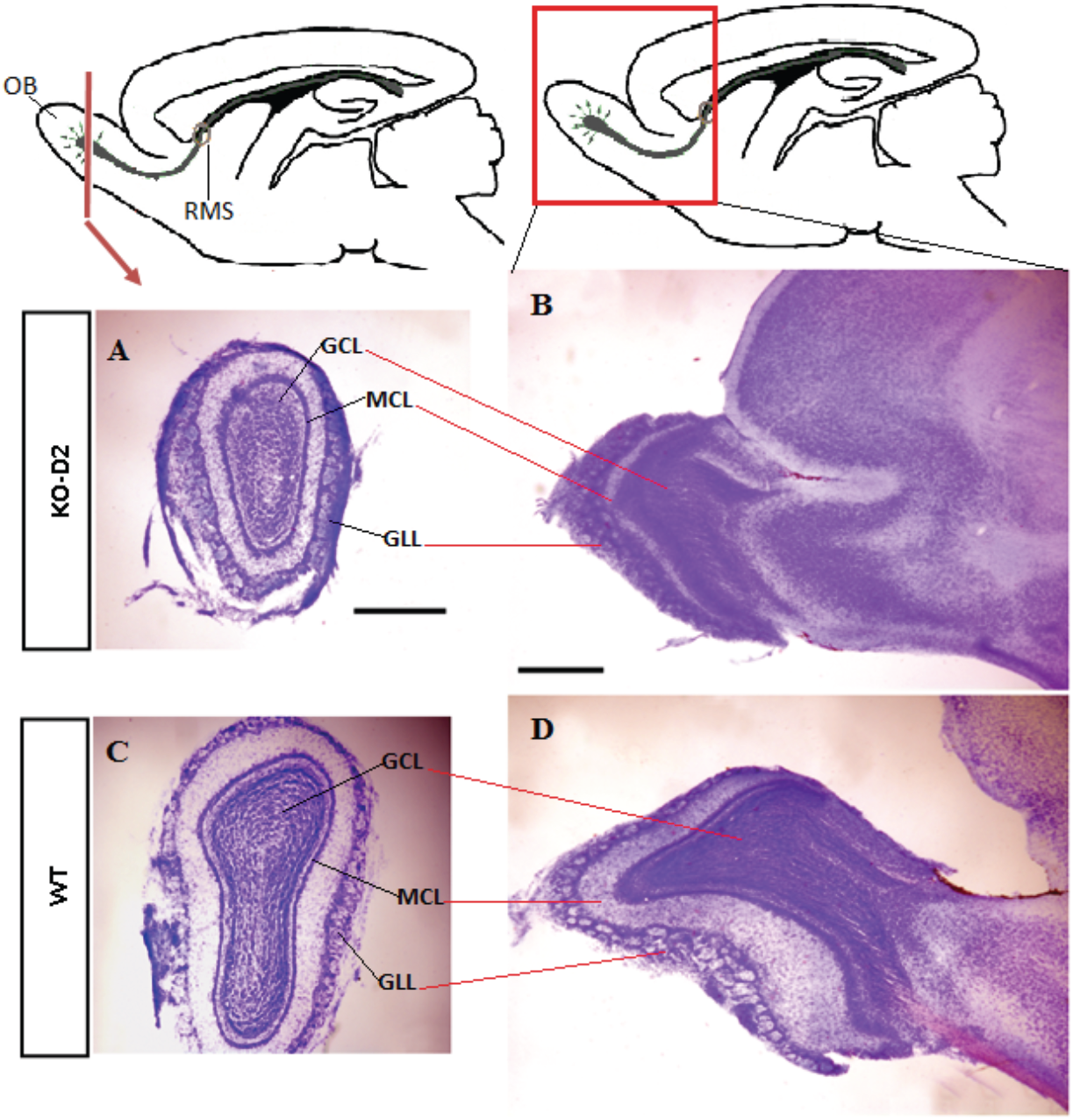
Cyclin D2-KO mice show altered olfactory bulb morphology. Olfactory bulb sections in the coronal plane (A and C) and in the sagittal plane (B and D) from the adult mice (P90). Top panel: cD2-KO mice. Bottom panel: cD2-WT mice. Histological analysis of the olfactory bulb morphology revealed that although all the layers are preserved in the cD2-KO mice, their thickness is altered and overall size is reduced. GCL - granule cell layer, MCL - mitral cell layer, GLL – glomerular layer. Scale bar 500 μm.

These observations were additionally in the support for investigation whether migrating neuroblasts within the rostral migratory stream (RMS) originating from the subventricular zone and present directly in the olfactory bulb have changed proliferative properties. Using EdU birthdate mapping, after a series of EdU injections to adult mice, we determined using epifluorescence microscopy that no EdU(+) cells were detected in the RMS within olfactory bulb in the cD2-KO mice, while WT mice showed frequent dividing events (Fig. 6). To confirm these observations, we processed olfactory bulb tissue for immunohistochemical detection of alternative endogenous marker of Ki67 protein, characteristic only for cells during all stages of mitotic division, and not present during G0 (interphase). Careful microscopic analysis confirmed that RMS within the olfactory bulb and their other areas present severely defective cell proliferation in cD2-KOs comparing to WT mice (Fig. 7). Performing visualization of neuroblasts with immunospecific antibody against doublecortin, we noticed that migration is changed and most of neurobladsts are arrested at the external plexiform layer, and migration of their subpopulation towards glomeruli is disrupted.

**Fig.6.**
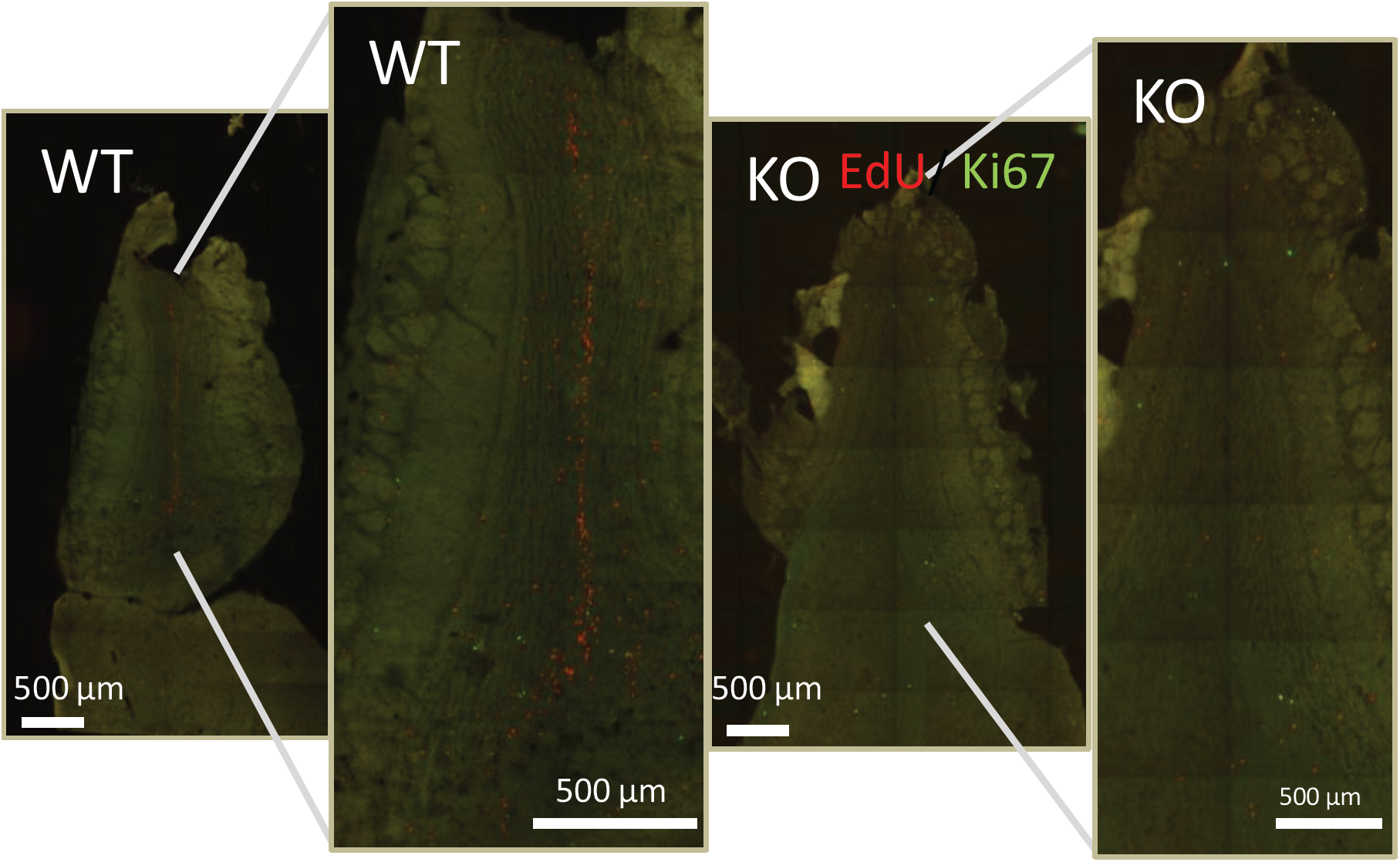
Attenuated proliferation activity and/or cellular migration of in cyclin D2-KO mice. Detection of EdU (red) and immunostaining for Ki67 (green) - proliferation markers, in the horizontal sections of the olfactory bulbs from cD2-WT (A) and cD2-KO (B) mice. Cyclin D2 deficiency markedly reduced the number of actively proliferating cells: EdU(+) and Ki67(+) profiles or migrating cells - EdU(+) profiles compared to cD2-WT animals (enlarged in A and B).

**Fig.7.**
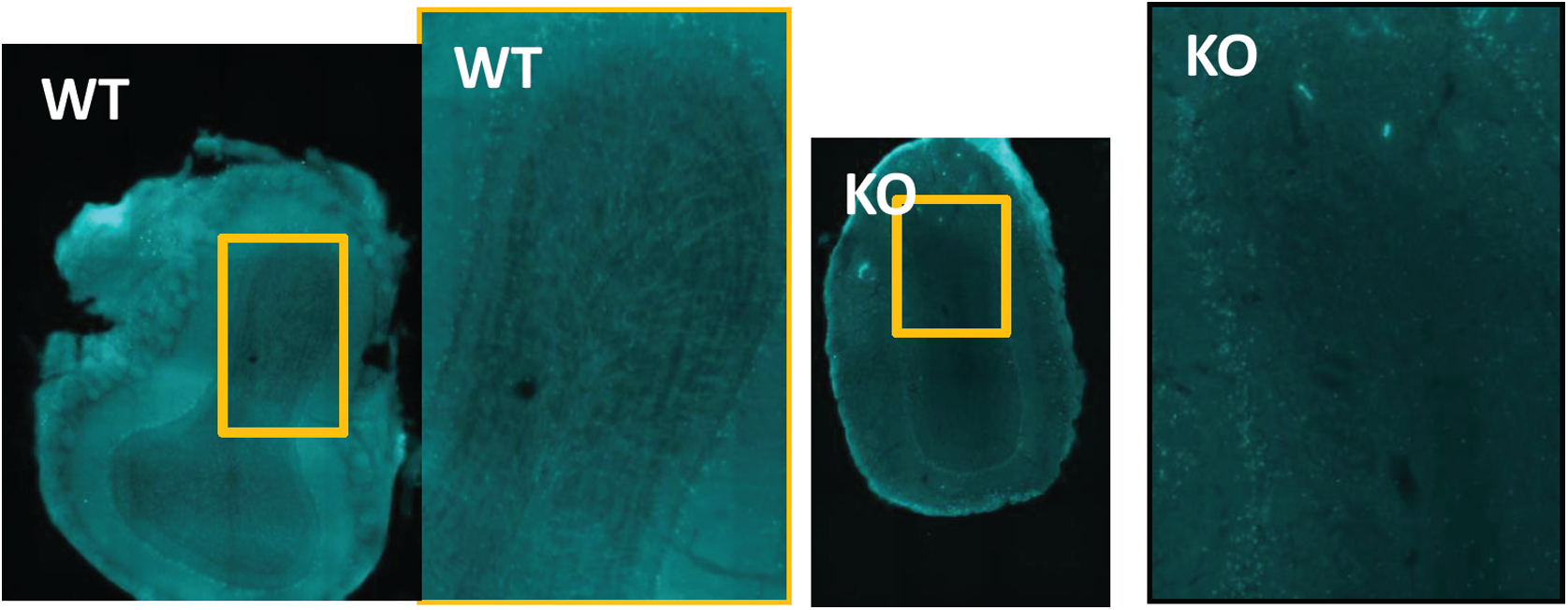
Cyclin D2-KO mice show reduced number of neuroblasts in the olfactory bulb. Immunostaining for doublecortin (Dcx) in the coronal sections of the olfactory bulbs from cD2-WT (A) and cD2-KO (B) mice. Microphotographs show reduced number of Dcx(+) neuroblasts in the center of the olfactory bulb (granule cell layer) in cD2-KO (B) compared to the cD2-WT animals (A).

In order to investigate interneuron subpopulation changes in the olfactory bulb as direct consequences of cumulative effects of changed proliferation in the SVZ, changed distribution of proliferating and migrating cells in the RMS and the olfactory bulb. We have employed immunehistochemical approach of identification of calretinin and calbindin interneuron subclasses, as well as dopaminergic interneuron, the latter with identification of tyrosine hydroxylase, a key enzyme in dopamine synthesis. We demonstrated that the number of calbindin-positive interneurons was decreased in the periglomerular layer. The number and distribution of calretinin and dopaminergic neurons seems to remain unchanged in cD2-KO mice (Fig. 8).

**Fig.8.**
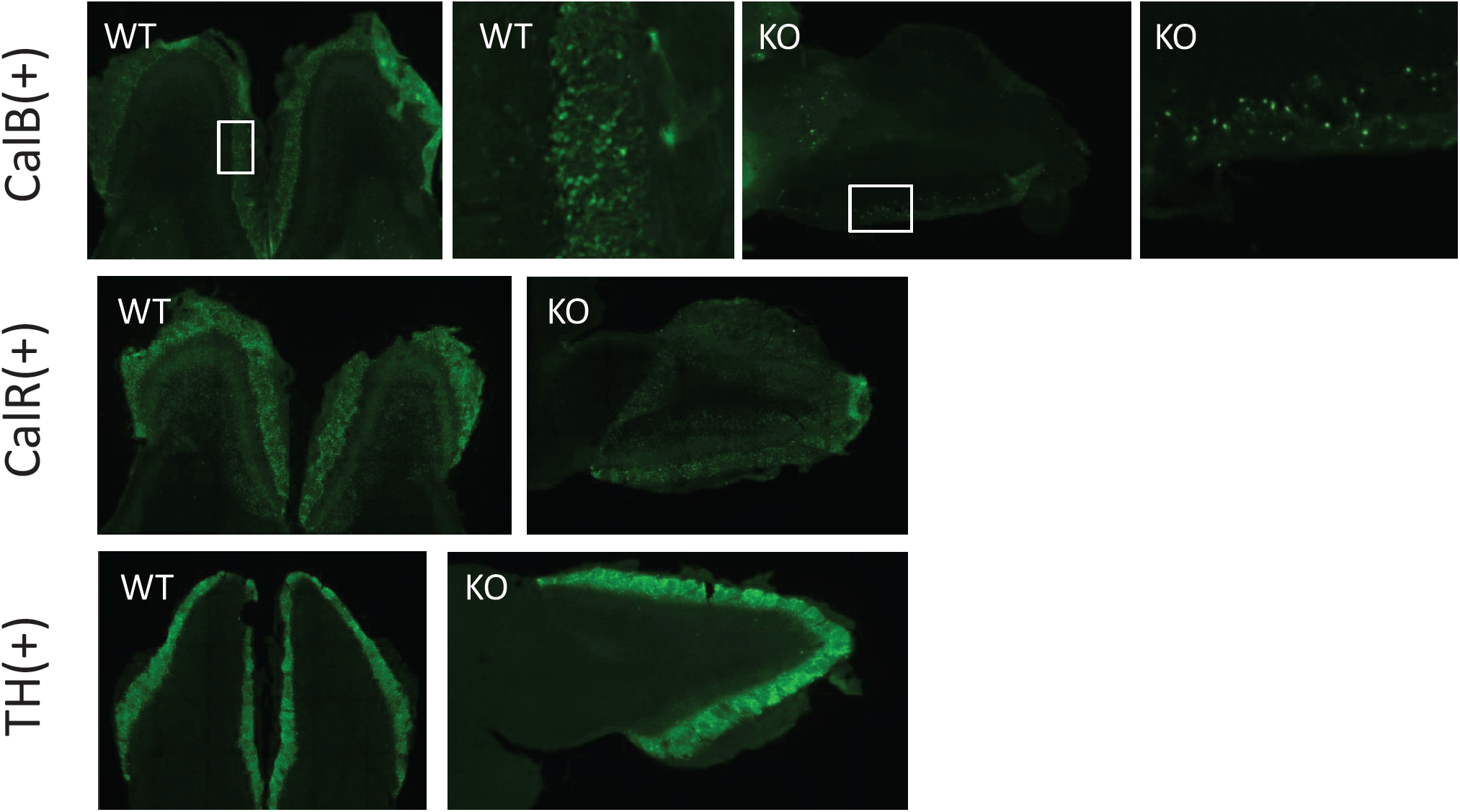
Olfactory bulbs of cD-KO mice are depleted of calbindin interneurons. Identification of the olfactory bulb interneurons with immunostaining for calbindin (A), calretinin (B), and tyrosine hydroxylase (C) from cD2-WT and cD2-KO mice. Cyclin D2 deficiency results in downregulation of CalB(+) interneurons in the glomerular layer of the olfactory bulb while there is no visible difference in CalR(+) or TH(+) interneurons compared to cD2-WT animals. CalB - calbindin, CalR - calretinin, TH - tyrosine hydroxylase.

## SUMMARY OF THE MAIN FINDINGS

In our experiments, we observed a general decrease in proliferation activity in most subregions of the SVZ niche in cyclin D2 knockout mice. Our results stay in line with previous reports from cD2-KO mice, documenting downregulation of proliferating cells in the hippocampus. Depletion of actively proliferating cells was not caused by upregulation of apoptosis as shown by immunodetection for active caspase 3 and using TUNEL method. Preliminary phenotypic analysis of the SVZ progenitors revealed that inactivation of cyclin D2 expression in adult mice affects the number of B1 active (activated precursors) and C type cells (intermediate progenitor cells) in the SVZ. This indicates an interruption in the transition from quiescent B1 cell stage to their activated form (active B1 cells). The most profound reduction in cell number was observed for neuroblasts, which were largely depleted in the SVZ and olfactory bulb of cyclin D2-deficient mice. The latter result may also indicate compromised migration of neuroblasts towards and within the olfactory bulb. Changes in proliferation activity of SVZ B1 precursors and intermediate progenitors after inactivation of cyclin D2 resulted finally in the altered composition of the olfactory bulb. We confirmed the overall decrease in size and shrinkage of cellular layers. The most pronounced observation, however, was the depletion of calbindin interneurons in the olfactory bulb glomerular layer of the cyclin D2 deficient mice.

## ACKNOWLEDGMENTS

The authors wish to express many thanks to people supporting our work: Aneta Schaap-Oziemlak, Agata Kania and Piotr Rogujski for help in maintaining cD2 mouse colony, Malgorzata Wieteska and Bartosz Zglinicki for help in tissue genotyping, and many others not listed for their support. We thank for funding this study to the National Science Center of Poland UMO-2012/07/B/NZ4/01733).

## STATEMENTS

The authors declare no competing interests.

